# Synthesis of cell-laden alginate microgels with tunable compositions based on microfluidic pico-injection technique

**DOI:** 10.1101/2022.01.09.475570

**Authors:** Yizhe Zhang, Angelo Mao, David J. Mooney, David A. Weitz

## Abstract

We report a microfluidic pico-injection-based approach for reliably generating monodisperse cell-laden alginate microgels whose composition can be tuned *in situ* through modulation of the cross-linker concentration. Separating the gelation from emulsification allows for a better control over the microgel size with a microfluidic drop-maker, and an instant adjustment of the microgel composition with a pico-injector.

## Introduction

Previous studies have revealed that as a major component of multicellular organisms, extracellular matrix (ECM) not only provides necessary mechanical and adhesive support to the cells, but also regulates various cell behaviors through facilitating the transmission of biochemical cues [1–6]. Systematic investigations on the interactions between the cell behaviors and its ECM can not only improve our understandings on the mechanisms underlying intricate cellular activities, but also contribute to the development of tissue engineering strategies for effective cell therapies [7–13]. Despite the vast advancement made in in-vivo studies, due to the complex and dynamic nature of physiological environment, in-vitro approaches are still widely applied in exploring some complex interplay between cells and their ECM, where cells are cultured and perturbed in bio-mimic scaffolds that are highly recapitulating their physiological microenvironment yet remain great flexibility in manipulatable components to decouple the entangled factors.

Lots of efforts have been made over the past two decades in developing suitable ECM scaffolds for in-vitro fundamental studies or tissue engineering applications [14–21]. Among many promising ECM materials, alginate hydrogel has been widely studied because of its good biocompatibility as a naturally derived ECM, and appealing structural properties that are amenable to chemical functionalization [22–33]. Alginate is the linear polysaccharide composed of repeating monomer units: β-D-mannuronic acid (M) and α-L-guluronic acid (G). Exposure to the divalent cation such as Ca^2+^ initiates the gelation process that crosslinks G blocks and forms the 3-D hydrated gel network. The considerable amount of hydroxyl and carboxyl sites on the polymer chain enables convenient chemical modifications to the hydrogel for desirable mechanical property and biodegradability. As the cell-laden scaffold, in most studies, the fabricated gels need to satisfy the requirement of long-term cell culture; therefore, it is ideal to reduce the size of the synthesized ECMs in order to minimize the diffusion barriers of nutrients, oxygen and biochemical cues. However, most of the reported external alginate microgel synthesis approaches lack the precise size control [34–41], whereas the large pore size and disturbed pH level from internal gelation strategies might interfere with cellular studies [42–49].

Based on the microfluidic pico-injector technique developed in our group, we designed a microfluidic two-step approach for alginate microgel fabrication, where drop-making process and gelation process were separated apart. Unlike the co-flow strategy typically used in gel bead synthesis on the microfluidic platform, our pico-injector employs the micron-sized injector nozzle to effectively limit the contact area of the alginate solution and the cross-linker flow, and therefore greatly suppresses the channel clogging at the fluid interface due to the rapid gelation, making the synthesis of alginate microgel more controllable, reliable and efficient on the microfluidic high-throughput platform. Moreover, no chemical addition or pH changes in our two-step approach ensures the fabricated microgel less invasive and thus more suitable for the long-term cell culture.

## Materials and Methods

### Alginate solution preparation

Alginate (Protanol LF 20/40; FMC Technologies) was functionalized with RGD peptide GGGGRGDSP of which 10% was Fluorescein isothiocyanate (FITC)-labeled for a uniform fluorescence under the confocal microscope (Peptides International) using standard carbodiimide chemistry as reported [50]. Briefly, for the DS (Degree of Substitution) 5 modification, 1 g of sodium alginate was dissolved to 1% (w/v) in a 0.1 M MES (Sigma) and 0.3 M sodium chloride (Sigma) buffer solution at pH 6.5, followed by the addition of 68.5 mg of sulfo-NHS (Pierce), 121 mg of N-(3-Dimethylaminopropyl)-N’-ethylcarbodiimide ( EDC) (Sigma) and 28 mg of RGD peptide. The reaction was allowed to proceed for 20 h before being quenched by the addition of hydroxylamine (Sigma). The reaction mixtures were then dialyzed against a decreasing concentration of sodium chloride for 2-3 days to remove salts and any unbound peptide, followed by the de-coloring step with activated charcoal. Finally the alginate was sterile (0.22 μm) filtered, lyophilized and reconstituted in serum-free DMEM (Dulbecco’s Modified Eagles Medium; Life Technologies) for ionic cross-linking.

### Cell suspension preparation

Clonally derived murine mesenchymal stem cells (mMSCs) (D1s; ATCC) were cultured in DMEM supplemented with 13 mM HEPES (4-(2-hydroxyethyl)-1-piperazineethanesulfonic acid; Life Technologies), 2 mM Sodium Pyruvate (Life Technologies), 10% FBS (Fetal Bovine Serum; Life Technologies) and 1% penicillin/streptomycin (Life Technologies) in a 37°C, 5% CO_2_ environment. The culture medium was refreshed every three days. To prepare the single-cell suspension suitable for infusion into microfluidic devices, the cells were trypsinized (Trypsin; Life Technologies), centrifuged at 1400 rpm for 5 min, and re-suspended into Dulbecco’s Phosphate Buffered Saline (dPBS; Life Technologies). The PBS wash was repeated a second time to remove unbound proteins. Cells were then re-suspended in serum-free DMEM/HEPES/Sodium Pyruvate for the final concentration of about 2 × 10^7^ per ml.

### Microfluidic device fabrication

Microfluidic devices were fabricated by patterning channels in polydimethylsiloxane (PDMS) using conventional soft lithography methods [51]. Briefly, for 15 μm drop-maker and 25 μm pico-injector that were used in our experiments, SU8-3015 and SU8-3025 photoresists (MicroChem Corp.) were spin-coated onto the 3” silicon wafers respectively and patterned by UV exposure through a photolithography mask. After baking and development with SU-8 developer (propylene glycol methyl ether acetate; MicroChem Corp.), the 15 μm and 25 μm tall positive masters of the devices were formed on the silicon wafers. Then a 10:1 (w/w) mixture of Sylgard 184 silicone elastomer and curing agent (Dow Corning Corp), degassed under vacuum, was poured onto the masters and cured at 65 °C for 2 hours. Afterwards, the structured PDMS replicas were peeled from the masters and inlet/outlet ports were punched out of the PDMS with a 0.75 mm-diameter biopsy punch (Harris Unicore). The PDMS replicas were then washed with isopropanol, dried with pressurized air, and bonded to 50 × 75 mm glass slides (VWR) using oxygen plasma treatment to form the devices.

For the pico-injector, to fabricate the electrodes into the device, a 0.1 M solution of 3-mercaptotrimethoxysilane (Gelest) in acetonitrile (99.8%; Sigma) was flushed through the electrode channels and blown out with pressurized air. A low melting point solder (Indalloy 19 (52 In, 32.5 Bi, 16.5 Sn) 0.020” diameter wire; Indium Corp.) was introduced into the electrode channels at 80 °C, followed by an eight-pin terminal block with male pins (DigiKey) glued with Loctite 352 (Henkel) to the surface of the device for strain relief. The solid electrodes in the shape of the channels were then formed when the device was cooled to the room temperature. Electrical contacts were made with alligator clips connected to a high voltage amplifier (Trek) and the function generator from the FPGA (field-programmable gate array) card running on the custom LabView program (National Instruments).

To enable the formation of aqueous-in-oil emulsions, the microfluidic channels were treated hydrophobic by flushing Aquapel (PPG Industries) into the device channels and immediately drying with pressurized air. To stabilize the drops against coalescence, we used EA surfactant donated by RainDance Technologies. The surfactant was dissolved in the fluorinated carrier oil Novec HFE-7500 (3M) at a concentration of 1.8% (w/w).

### Microfluidic operation

All the microfluidic devices, connecting tubing (Fisher Scientific), plastic syringes (BD), needles (BD), collecting tubes (Eppendorf), pipette tips (VWR) were sterilized with UV illumination. A flow of 70% ethanol through the tubing, syringes and needles was applied before each round of experiment to prevent the bacterial contamination. PhD 2000 syringe pumps (Harvard Apparatus, Inc.) were used to infuse the fluids into the device.

To produce the cell-encapsulated alginate drops, freshly prepared D1 mMSC suspension was infused into the inner channel of the 15 μm drop-maker at a flow rate of 60 μl/h. The 2.5% (w/v) alginate solution was co-flowed in the middle channel at the flow rate of 60 μl/h to form the cell-alginate mixture at the first junction of the device before sheered into the droplet at the second junction by the fluorinated oil HFE 7500 that contained EA surfactant in the outer channel with the flow rate of 240 μl/h.

The drops coming out of the drop-maker were collected into a 1 ml plastic syringe, re-injected into a 25 μm pico-injector at 80 μl/h and spaced out evenly by the EA-containing oil HFE 7500 flowing in the outer channel at 800 μl/h. 100 mM calcium chloride solution was injected from the pico-injecting side channel into the drops at a flow rate of 20 μl/h. Under the rupturing effect [52] of an electric field (80~90 Vpp, 30 kHz), the alginate drop surface was destabilized and got fused with the pico-injecting calcium chloride to initiate the cross-linking. The precursors and the cross-linkers in the pico-injected drops were then fully mixed from advection in the sinusoidal channel adjacent to the pico-injecting junction, and the serpentine channel downstream allowed for the sufficient gelation before the drops were creaming in the collecting tube.

### Gel exaction from the oil phase

After the gelation process, the collected emulsion was washed with 20% (v/v) Perfluorooctane (PFO; Sigma) in HFE 7500 solution to remove the oil phase and the cell-gel pellet was re-suspended in the serum-free DMEM supplemented with HEPES and Sodium Pyruvate for long-term culture.

### Cytotoxicity test on ECM-culturing cells

Trypan blue based exclusion assay was employed to assess the cell viability cultured on the in-vitro ECM. A 1:1 (v/v) mixture of the cell-gel suspension and the sterile (0.22 μm) filtered trypan blue solution (0.4% trypan blue in 0.81% sodium chloride and 0.06% dibasic potassium phosphate; Life Technologies) was prepared and incubated for 2 min at room temperature for viable/non-viable cell imaging under the phase contrast microscope. In some imaging, cell membrane dye PKH26 (Sigma) was employed for a better identification of the cells with regard to the scaffold gel beads.

## Results and Discussion

Fig. 1 describes the two-step synthesis process, with the drop-maker (Fig. 1a) producing aqueous alginate drops, followed by the pico-injector (Fig. 1b) serving as the gelation platform. The ultra-small size of the injecting nozzle reduces the possibility of clogging at the interface of the uncross-linked flow and cross-linker flow, whereas the evenly spaced single-file flow of the alginate drops allows a tight control of the cross-linker addition for an improved uniformity on the microgel structure. The incorporation of the sinusoidal design in the downstream of the T-junction enables an efficient mixing of the calcium ions and alginate solution in the droplet, and the subsequent serpentine channel allows for the sufficient gelation before the drops cream in the collection tube. Typical drop-making process is demonstrated in Fig. 1c, where encapsulated mMSCs are highlighted with the red arrows. Fig. 1d is the time-sequence snapshots from a pico-injecting process captured by the fast camera (Phantom V7.3; VisionResearch). The entire cross-linker-adding process was completed within 1.25 millisecond, which gives a kilo Hertz production rate. By controlling the cell density infused into the drop-maker, we can control the cell number in each droplet. Through adjusting the flow rates in the pico-injecting process, we can control the ratio of the cross-linker to the alginate solution in each gel bead for controllable mechanical structure. And benefited from the clean gelation process, the pico-injector devices could usually be reused for multiple rounds of experiments, which greatly reduced the cost in time and money.

**Fig. 1.**
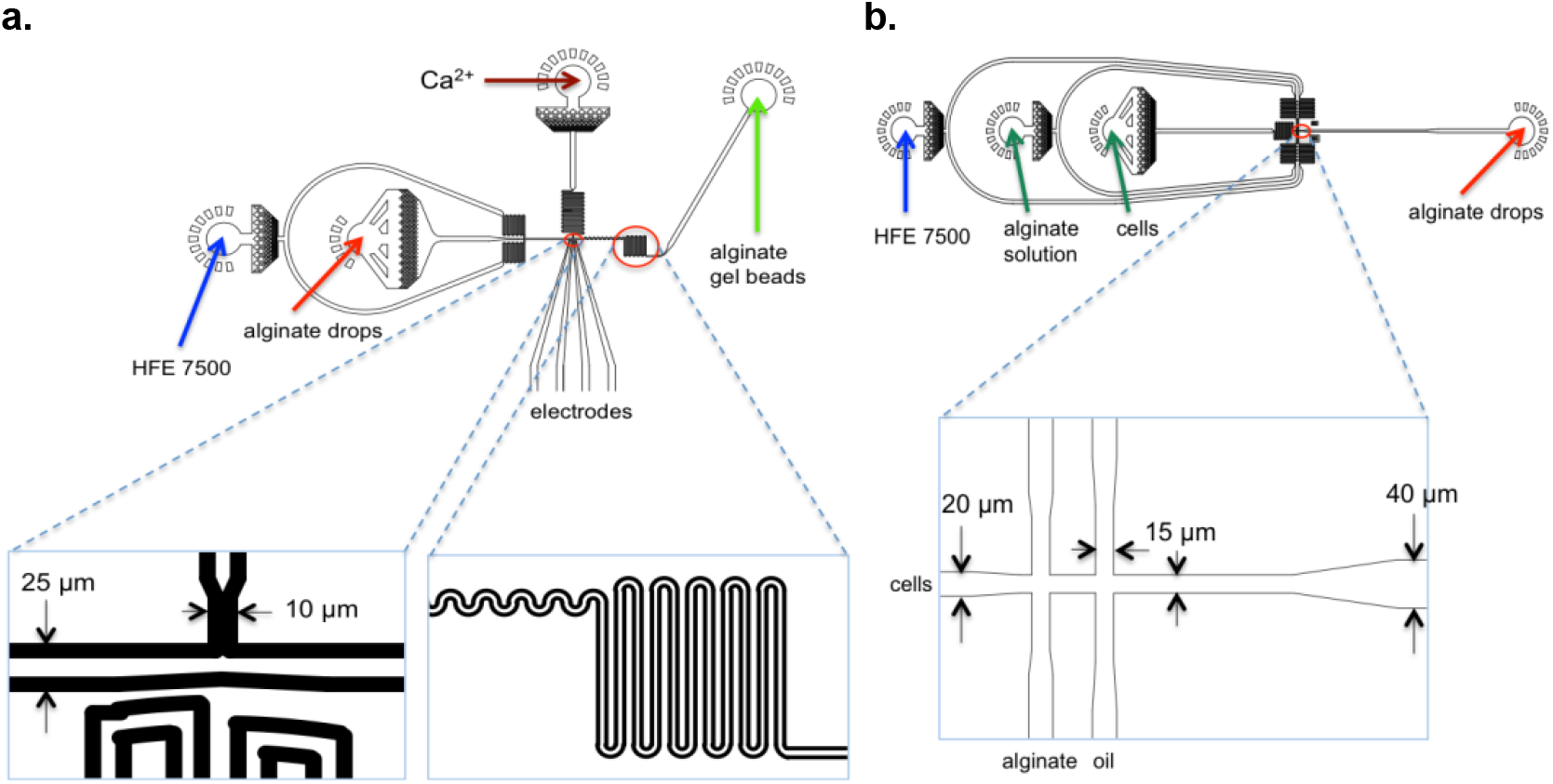
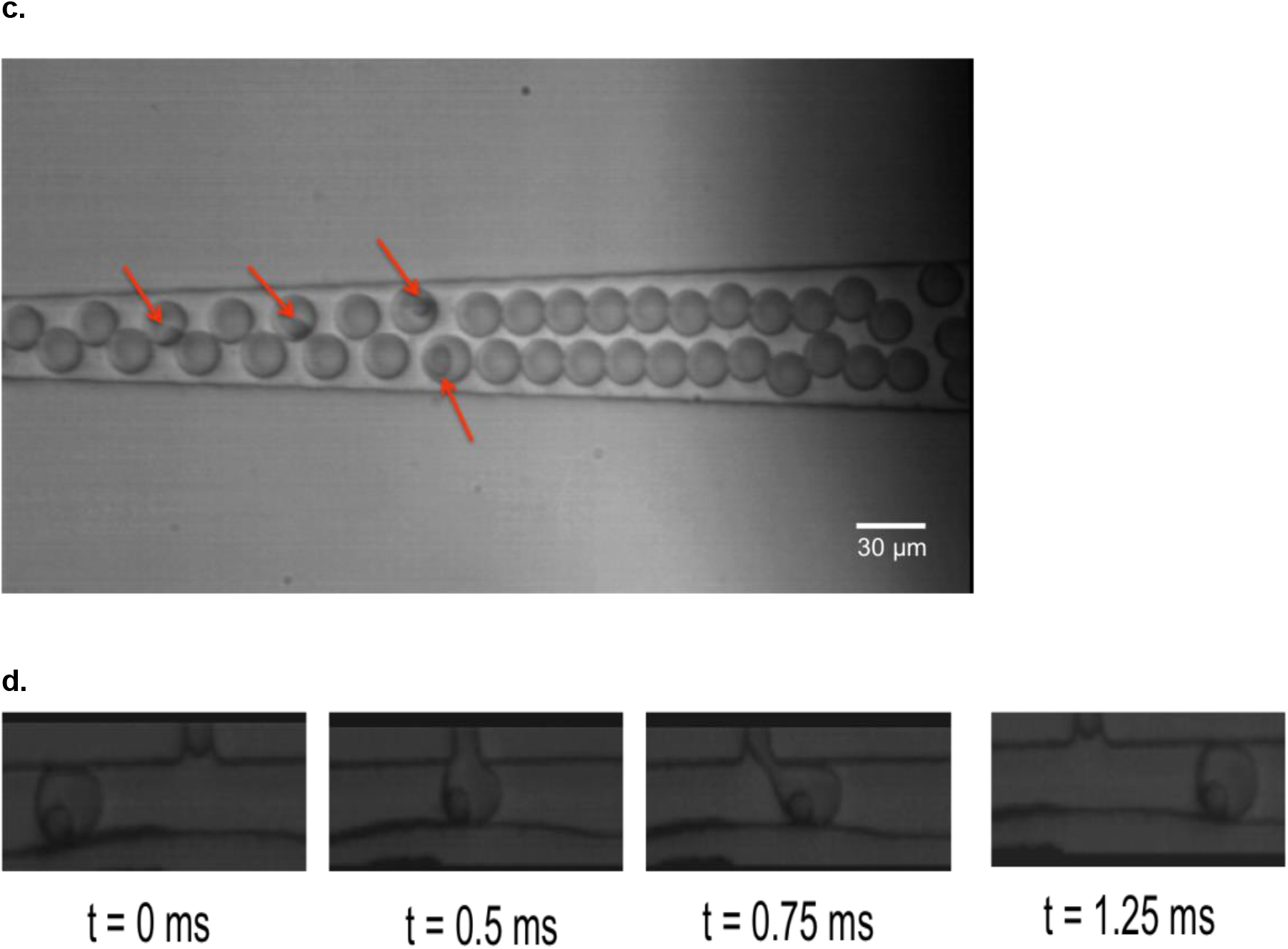
Two-step synthesis of alginate microgel with microfluidic devices. **(a)** Schematic of the drop-maker device: Alginate solution and mesenchymal stem cells were co-flowed into the aqueous channel from left to right. Fluorinated oil HFE 7500 with EA surfactant was flowed into the oil channel and sheered the aqueous fluid into ~25 μm drops at the junction downstream of the serpentine resistant channel. **(b)** Schematic of the pico-injector device: Alginate drops were injected into the inner channel and spaced out by the carrier oil HFE 7500 before flowing right to the T-shaped pico-injecting nozzle. The characteristic design downstream to the injecting junction (highlighted in the red circle) is intended for improved mixing and sufficient gelation on chip. **(c)** Snapshot of the movie depicting the cell encapsulation into alginate drops. The encapsulated mesenchymal stem cells were marked with the red arrows in the picture. **(d)** Time-sequence snapshots from a pico-injecting movie. In the representative snapshots, the cell-laden alginate drop was injected with calcium chloride from the perpendicular pico-injecting channel for gelation initiation. The fusion of the injecting cross-linker solution into the alginate aqueous drop was accomplished upon the activation of the electric field applied at the injecting area. The injection step was finished within 1.25 ms.

We first synthesized empty alginate gel beads to obtain the optimum gelation conditions. We found that both the calcium concentration and its injected rate affected the gelation process. To remove the incomplete gelation products, freshly produced gel beads were washed out of the carrier oil into the cell medium. For 1% alginate, only when the concentration of the calcium chloride reached above 100 mM in the injecting channel (20 mM in the droplet), the maximum number of completely gelled beads was obtained, as indicated in Fig. 2a. And the injecting rate of the calcium chloride had a great effect on the homogeneity of the meshwork in the gel bead. For a fixed calcium concentration, the increased injecting rate to the alginate drop resulted in an increased homogeneity in the spatial distribution of the cross-linking (Fig. 2b). However, the increase in the injecting rate lead to the gel size increase, and in a confined channel, resulting in a polarized shape. For a trade-off between the internal homogeneity and the isotropic shape, we chose a calcium concentration of 20 mM in drops injected at a 20% rate (v/v) of the final droplet volume.

**Fig. 2.**
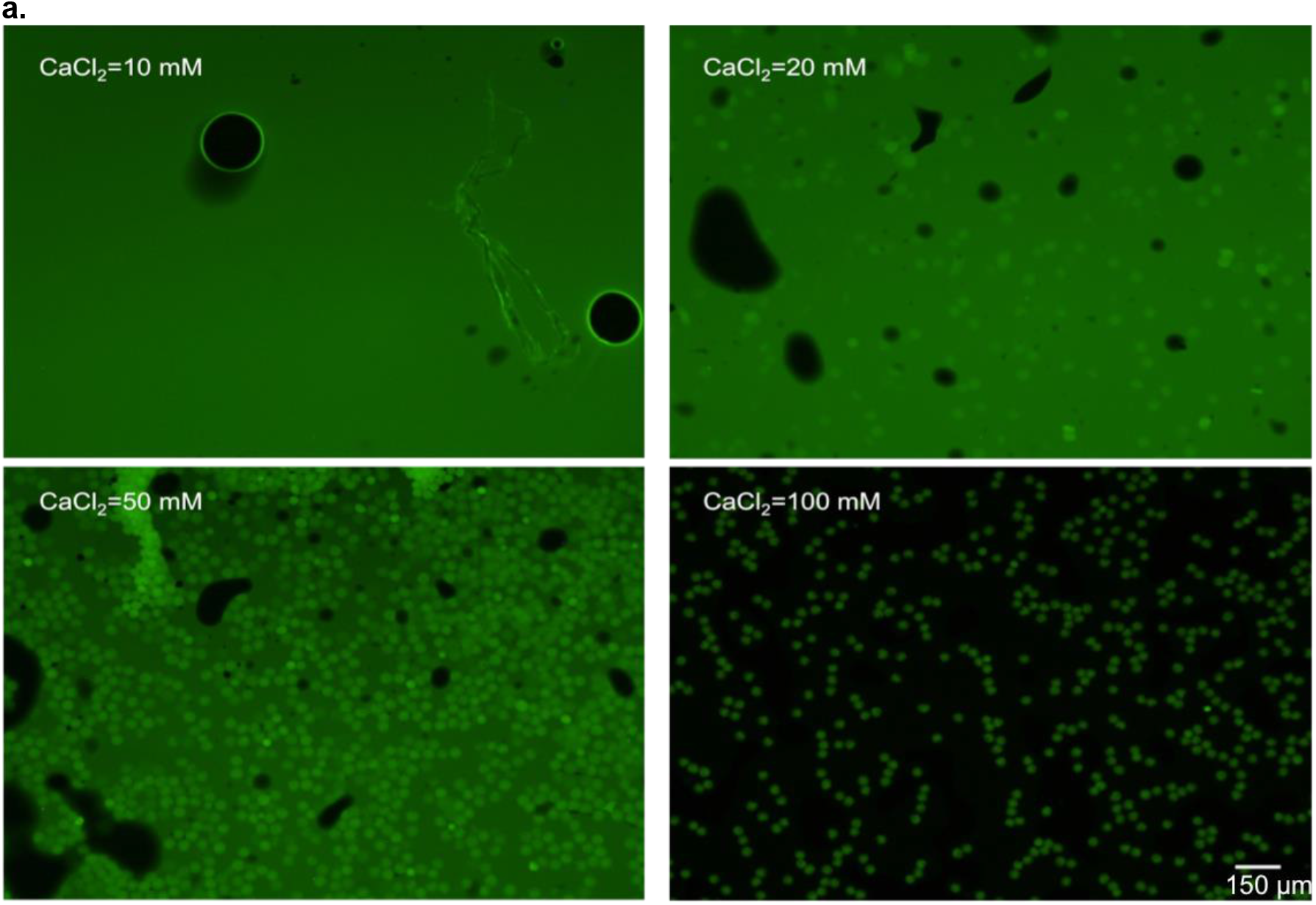
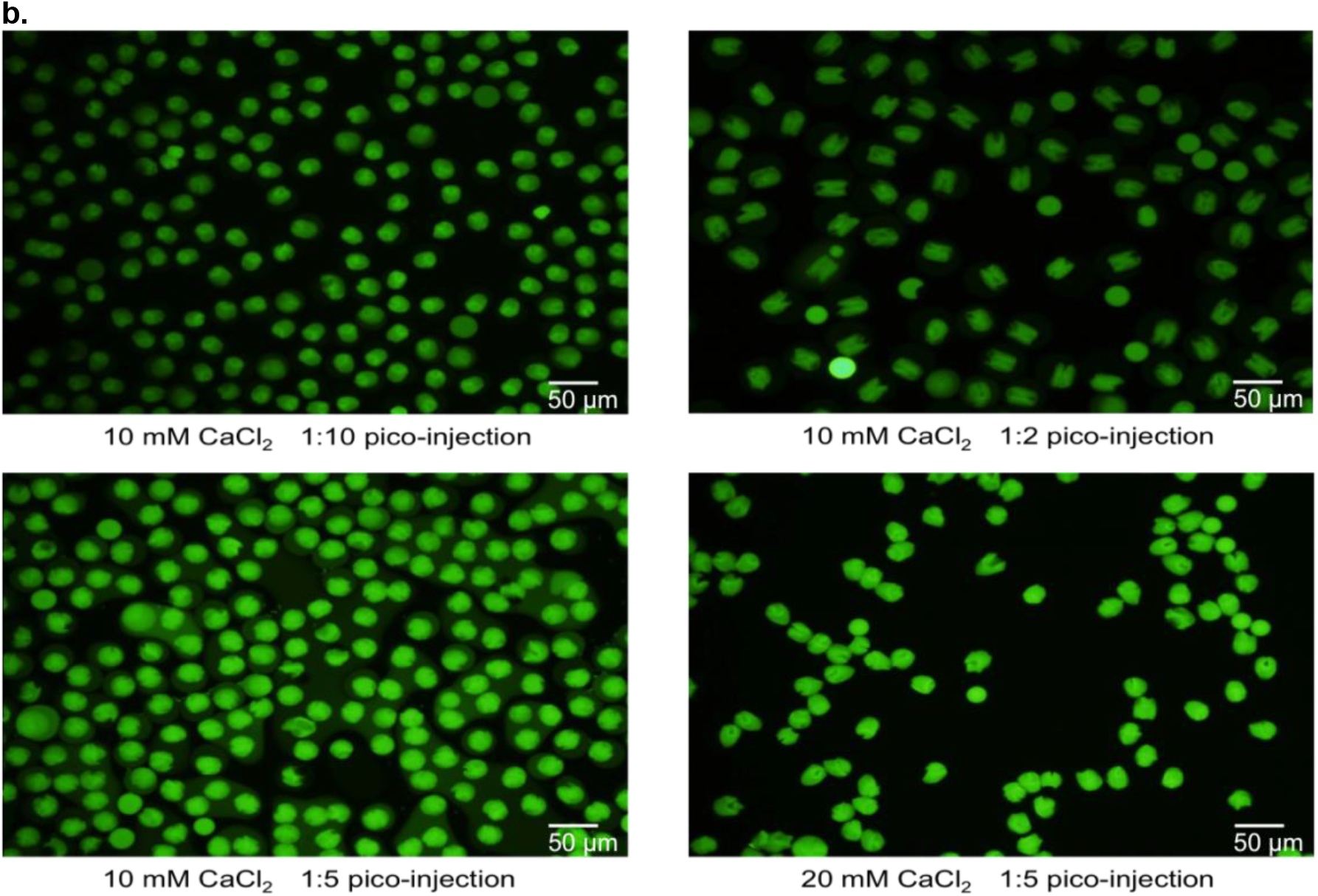
Optimization of the gelation conditions. (a) Fluorescence microscopy of gel beads formed from different Ca^2+^ concentrations. The displayed values are the concentrations in the pico-injecting channel. (b) Fluorescence microscopy of gel beads formed with different Ca^2+^ injecting rates. The ratios are the volume ratio of the injecting fluid to the injected alginate drop. The CaCl_2_ concentrations are the expected concentration in the injected drop. A fraction of alginate chains was pre-functionalized with green fluorophore FITC for clarity. The gel beads in (a) and (b) were both washed from the carrier oil into the cell culture medium before imaging.

We then incorporated mMSCs into the gel beads by encapsulating them in the alginate drops. For necessary cell adherence, alginate was modified with the cell-binding peptide arginine-glycine-aspartate (RGD). For easy optical identification, a fraction of the RGD peptides were also functionalized with green fluorophore Fluorescein isothiocyanate (FITC). Fig. 3a (top left) shows the fluorescence image captured right after the gel beads were formed. The beads that contained the cells are indicated in red circles. The examination on the Trypan blue-stained cells in the phase contrast microscopic images showed more than 90% viability of the gel-encapsulated cells, as represented in the top right picture. Similar cell encapsulation experiment was also performed with the red membrane dye PKH26-stained cells and the fluorescence images were taken right after the gel beads formation. As clearly displayed in Fig. 1a (bottom), quite a number of cells were encapsulated in the ~40 μm alginate gel scaffold, with the existence of both single- and double-occupancy. The observation that cells tended to reside at the edge of the gel bead might be associated with heterogeneous cross-linking within the gel beads, which is possibly the effect of the diffusion barrier formed by the fast cross-linking at the contact area of the alginate droplet and the injected fluid that obstructed the further diffusion of more calcium ions into the drop. It has been observed in other external gelation approaches too and integrating sodium chloride into the cross-linker calcium chloride was reported to be able to effectively suppress this phenomenon [53]. Nevertheless, the cells were successfully trapped in the 3D micro scaffold.

**Fig. 3.**
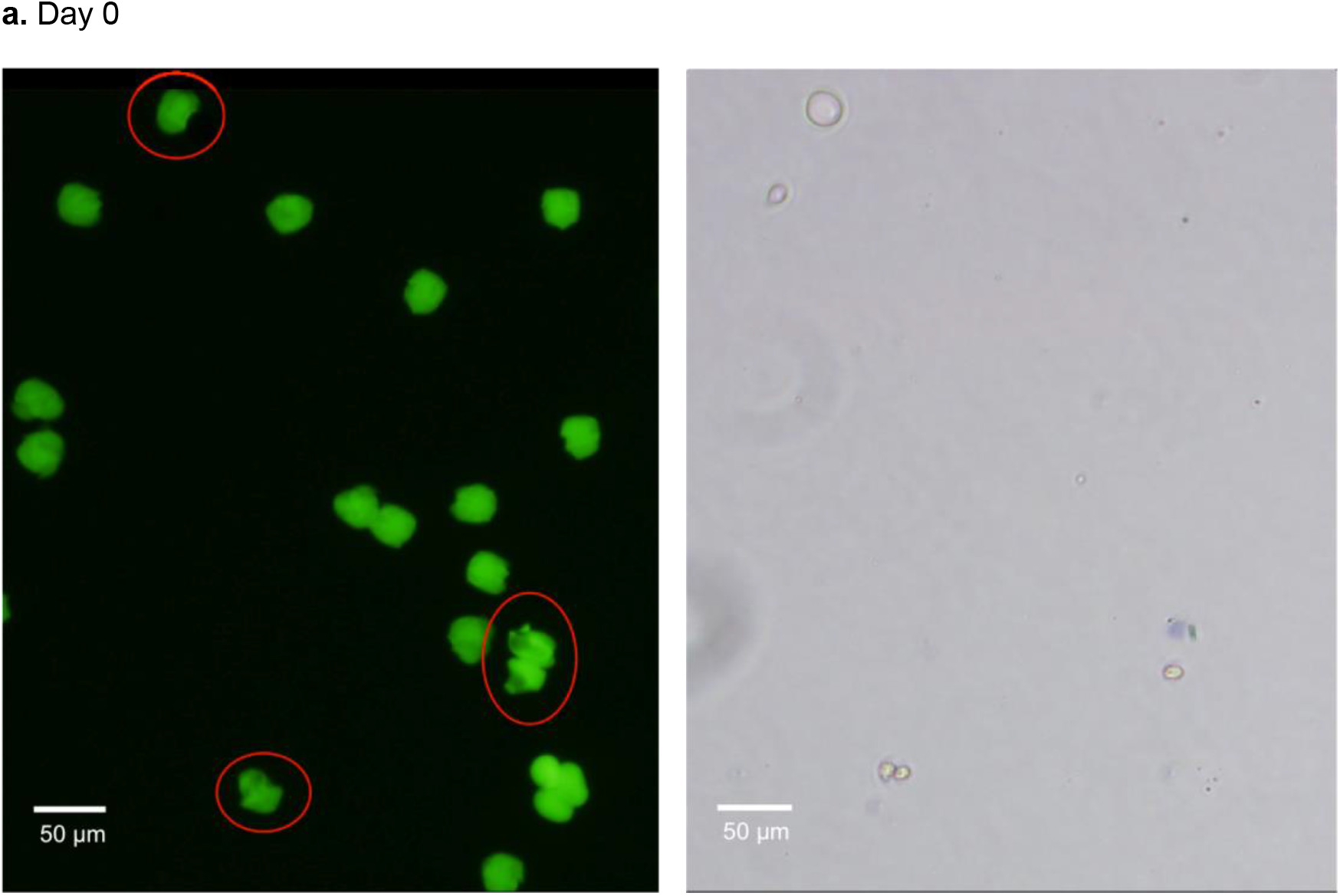
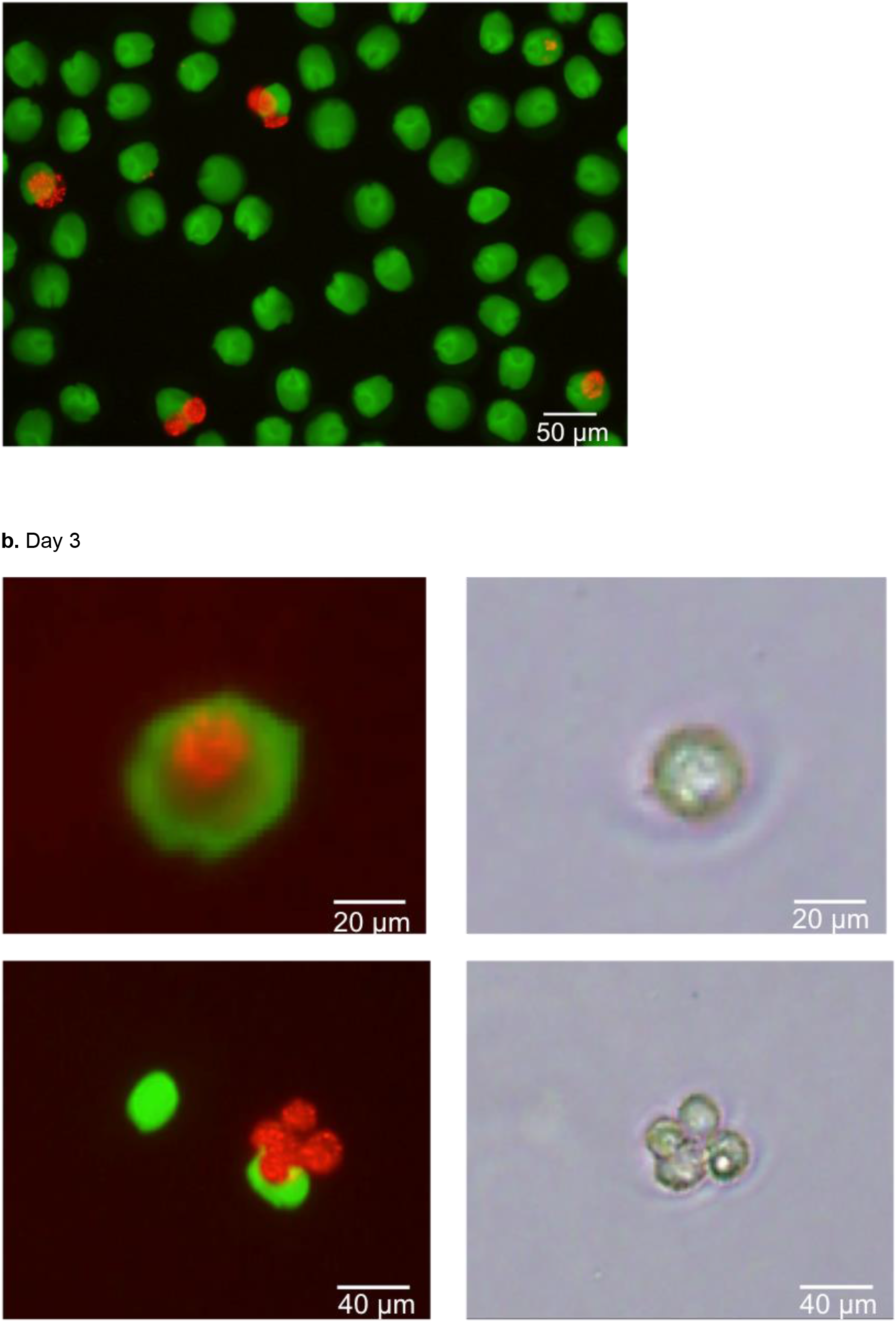
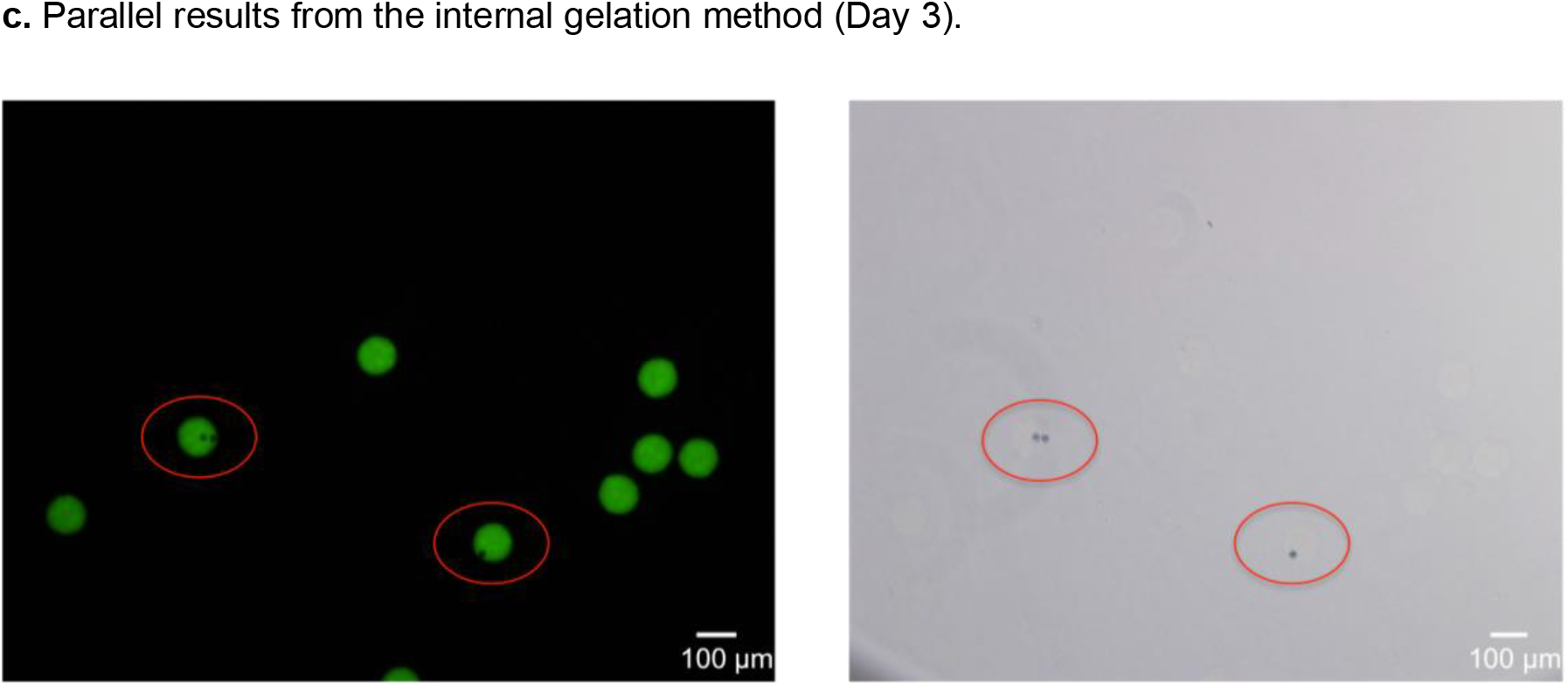
Cell viability measurement on the encapsulated mesenchymal stem cells. (a) Representative fluorescence (top left) and phase contrast (top right) micrographs of the cell-laden gel beads right after the gelation process. Red circles highlight the cell-laden gel beads. Trypan blue-based exclusion assay showed no dead cells in the examined field. The bottom picture is the overlay of fluorescence micrographs from the red and green channels, to offer a clear view of cells’ distribution in the gel. (b) Fluorescence (left; overlay from red and green channels) and phase contrast (right) micrographs of the live cells based on Trypan blue assay after 3-day culture in the gel beads. Cells were stained with the red membrane dye PKH26 for clarity. (c) Fluorescence (left) and phase contrast (right) micrographs of the cell-laden gel beads produced from the internal gelation method. Trypan blue assay was performed on Day 3. Red circles highlight the cell-laden gel beads.

We cultured these gel beads in the cell medium to examine the biocompatibility of the ECM synthesized with our microfluidic pico-injector-based method. On Day 3, a fraction of the beads was stained with Trypan blue and examined under confocal microscope and phase contrast microscope for cell viability assay. We got a nearly 86% of viability in all the trapped cells examined. Some cells even exhibited a proliferation tendency in the microscopic image (Fig. 3b), where a group of cells were physically associated with each other in the immediate vicinity of the gel bead. As a comparison, a parallel experiment was performed on the gel beads synthesized from the internal gelation method where alginate was encapsulated with calcium carbonate nanoparticles into the aqueous droplets surrounded by the 0.3% (v/v) acetic acid-incorporated oil phase. The viability assay showed no live cells present in the sampling gel beads on Day 3 (Fig. 3c). Although more data on the longer-term culture may be needed to assess the biocompatibility of the pico-injector approach by taking account of its effect on the cell proliferation and differentiation, the results we had here at least suggested that our pico-injector approach is promising as a less-invasive 3D ECM synthesis method compared to other internal gelation strategies.

## Conclusion

We have successfully synthesized alginate microgel that can be reliably produced down to 30 μm in size on a pico-injector-based microfluidic platform. It pushes the gel fabrication towards the lower size limit as ECM for mammalian cells, which is especially desirable for long-term in-vitro studies on isolated cell behaviors, such as the stem cell differentiation in the model stem cell niche. The cell viability assay performed on Day 3 suggested the mild invasiveness of this acid-free gelation approach on the mesenchymal stem cells that were seeded inside the synthesized scaffolds, which is critical for, for example, stem cell niche studies that require the integrity of the cells for lineage commitment observations under the regulation of the specific signals. In addition, the highly-scalable feature of this approach also enables the mass production of the cell-laden scaffolds for cell therapy in clinical applications.

By regulating the input cell density in the drop-maker and the flow rates in the pico-injector, we can also have a fine control over the encapsulated cell number and the gel composition. Furthermore, through a secondary injection, this pico-injector gelation platform can be easily adapted to produce multi-niches within one aqueous droplet for controllable studies of cell-cell interaction in the isolated microenvironment.

## Acknowledgements

This work was supported by the National Science Foundation (DMR-1310266) and the Harvard Materials Research Science and Engineering Center (DMR-1420570).

